# In vitro culture of bovine fibroblasts using select serum-free media supplemented with *Chlorella vulgaris* extract

**DOI:** 10.1101/2021.11.17.469028

**Authors:** Galileo Defendi-Cho, Timothy M. Gould

## Abstract

Standard cell culture practices require addition of animal-derived serum to culture media to achieve adequate cell growth. Typically, 5-10% by volume of fetal bovine serum (FBS) is used, which accounts for a vast majority of the cost of media while also imposing environmental and ethical concerns associated with the use of animal serum. Here we tested the efficacy of culturing cells by replacing serum in the media with algae extract and select additives. Using LC-MS, we compared molecular signatures of FBS to *Chlorella* algae extracts and identified NAD(H)/NADP(H) as common and relatively abundant features in their characteristic profiles. Bovine fibroblasts, cultured in serum-free media supplemented with *C. vulgaris* extract and just two growth factors plus insulin, showed significant growth with enhanced viability compared to control cells cultured without serum, albeit still lower than that of controls cultured with 10% FBS. Moreover, *C. vulgaris* extract enhanced cell viability beyond that of cells cultured with the two growth factors and insulin alone. These results suggest that key components in serum which are essential for cell growth may also be present in *C. vulgaris* extract, demonstrating that it may be used at least as a partial alternative to serum for cell culture applications.

## Introduction

In 2013 it was estimated that raising livestock contributed roughly 14% to total greenhouse gas emissions, more than half of which was attributable to meat production (Gerber et al. 2013). Cell cultured meat has the potential to offer substantial environmental and social advantages over traditional agriculture practices using livestock (Tuomisto and Teixeira De Mattos 2011; Ladak and Anthis 2021), yet analysis on the economics of cultured versus conventional meat production shows high cost barriers, particularly those associated with the nutrients required (Humbird 2021). Cell culture media is commonly supplemented with fetal bovine serum (FBS) because animal serum promotes cell growth (Penttinen and Saxén 1959). Based on current commercial prices, the cost of culture-grade FBS is estimated at around $1.20 per mL and that of other media components at $0.04 per mL; thus, for typical complete growth media containing 10% FBS, most of its total cost is attributed to FBS, by roughly three-fold. Therefore, low cost and environmentally favorable alternatives to serum are attractive for cell culture applications, particularly for the cultured meat industry.

Because animal-derived serums like FBS are very complex, including thousands of different molecules that can be highly varied in their relative abundance due to many dynamic physiological factors, it is unlikely that synthetic approaches to serum alternatives will prove cost effective if technically feasible. Researchers have studied the culture of cells in serum-free conditions for many decades with varying degrees of success, yet most cell culture practices continue using FBS. Following studies which identified key components necessary to culture cells in the absence of serum (Murakami and Masui 1980; Chen et al. 2011), there are now chemically-defined serum-free media formulations that are commercially available (e.g., E8™).

However, the cost of these serum-free formulations is still quite high, primarily due to the protein growth factors required. Such formulations were developed mainly for the purpose of consistently culturing human induced pluripotent stem cells (iPSCs); they were not originally intended for cultured meat applications although this does typically involve culturing stem cells. Nevertheless, determining the minimal components and concentrations at which they are needed to achieve adequate and consistent cell growth remains an important undertaking for the cultured meat industry (Specht 2020).

We were intrigued by a recent study in which the authors reported growth effects of algae extract on mammalian cells (Ng et al. 2020). Ng and colleagues report that extract derived from *Chlorella vulgaris* promoted growth of Chinese hamster ovary (CHO) and mesenchymal stem cells (MSC) under certain conditions, some of which did not contain any serum. In another study, researchers reported high levels of glucose and amino acids present in various microalgae extracts, with particularly high levels of aspartate and glutamate detected in extract from *C. vulgaris* (Okamoto et al. 2020); these authors proceeded to show that they could use such algae extracts as a substitute to culture cells in media deficient in glucose and amino acids. More recently, this same group showed that *C. vulgaris* extract can be used as a replacement for DMEM in the culture of primary bovine myoblasts (Okamoto et al. 2022).

Here we tested the efficacy of culturing cells by replacing serum in the media with algae extract and found that we could achieve appreciable growth and viability of bovine fibroblasts using select serum-free media supplemented with *C. vulgaris* extract (CVE). We characterized molecular features of algae extracts, compared them to that of FBS, and found evidence suggesting that NAD(H)/NADP(H) are common and relatively abundant components. This study establishes how algae extract may be used to culture cells in serum-free conditions while providing further insight into essential components required for cell growth.

## Materials and Methods

### Chemicals and Reagents

Liquid chlorella growth factor (SKU 504) and *Chlorella vulgaris* extract capsules (SKU 583) were purchased from BioPure® Healing Products (Woodinville, WA). Pierce™ universal nuclease for cell lysis was purchased from Thermo Fisher Scientific Corporation (Waltham, MA). Recombinant human fibroblast growth factor (FGF-2, catalog no. 100-18B) and transforming growth factor beta (TGF-β, catalog no. 100-21) were purchased from PeproTech, Inc. (Cranbury, NJ). Bovine insulin, sodium L-ascorbate and selenium dioxide were from Sigma-Aldrich. Cell titer 96® aqueous one solution cell proliferation (MTS) assay (catalog no. G3580) was purchased from Promega Corporation (Madison, WI).

### Fractionation of Algae Extracts

Algae extracts termed *Chlorella vulgaris* extract (CVE) and chlorella growth factor (CGF) were derived from two different strains, *Chlorella vulgaris* (CVE) and *Chlorella pyrenoidosa* (CGF). CVE was obtained from capsules in dry powder form and dissolved directly in serum-free DMEM. CGF stock solution was purchased as a concentrated liquid hot-water extract from which all solutions were made on a percent volume basis by diluting CGF directly in serum-free DMEM. For fractionation experiments, crude solutions of 10 g/L CVE and 10% CGF were first prepared in serum-free DMEM. Crude CVE and CGF were then either sterile-filtered through 0.22 μm nylon filters or centrifuged at 10,000 x *g* for 10 minutes in a benchtop microcentrifuge. After centrifugation, supernatants were removed, and the pellets were resuspended directly in serum-free DMEM.

### LC-MS

Sample separation was performed by liquid chromatography (LC) using a UPLC BEH C18 column (2.1 × 50 mm, 1.7 μm) fitted to an Acquity UPLC system with PDA detector (Waters). Chromatographic profiles were acquired at wavelengths 200-400 nm. Injection volume was 7.5 μL after equilibration, and the column temperature was held at 30°C. Samples were eluted from the column using a gradient mobile phase that consisted of phase A (0.1% formic acid in water) and phase B (0.1% formic acid in acetonitrile). The linear gradient elution procedure was as follows: (step 1) 100% A from 0-3 minutes, (step 2) 100% to 60% A from 3-20 minutes, (step 3) 60% to 0% A from 20-21 minutes, (step 4) 0% A from 21-24 minutes, (step 5) 0% to 100% A from 24-25 minutes. The column flow rate was set at 0.2 mL/min from 0-21 minutes, increased to 0.6 mL/min from 21-24 minutes, and then returned to the initial rate from 24-25 minutes. Electrospray ionization mass spectrometry (ESI-MS) was performed using an Acquity QDa detector (Waters). Capillary and cone voltages were 0.8 kV and 10 V, respectively. MS measurements were collected in positive ion mode with a mass scan range from 100-1200 *m*/*z*.

### Cell Culture

The EBTr (NBL-4) cell line was purchased from ATCC (CCL-44™). Cells were routinely cultured at 37°C with 5% CO_2_ in complete Dulbecco’s Modified Eagle’s Medium (DMEM} supplemented with 10% FBS (Gibco), 100 U/mL penicillin and 100 μg/mL streptomycin (Thermo Fisher Scientific, catalog no. 15140148); serum-free DMEM included antibiotics but no FBS. Passaging was routinely done as follows: cells were first washed with 1X phosphate-buffered saline (PBS), then placed in an incubator with 0.25% Trypsin-EDTA solution (Gibco) for approximately 2 minutes immediately after which complete DMEM was added to quench trypsin activity; cells were collected and centrifuged at 300 × *g* for 5 minutes, and then the cell pellet was resuspended in fresh media. EBTr cells were passaged every 3-4 days at a 1:3 passage ratio. Excess cells were stored in liquid nitrogen at 1-2×10^6^ cells/mL in freezing media containing 90:10% FBS:DMSO. The number of passages for experiments was limited to less than twenty.

### Viability (MTS) Assays

EBTr cells were first harvested, counted, and resuspended at a density of 2.5×10^5^ cells/mL in serum-free DMEM. Cells were then seeded in clear, flat-bottom 96-well plates to achieve a target density of approximately 2×10^4^ cells (80 μL) per well. Seeded cells were incubated for 2-4 hours to allow sufficient time for cells to adhere to plates prior to experiments. For all viability experiments/assays, each condition corresponded to a single column in a 96-well plate. Test media conditions were achieved by delivering 20 μL of 5X test solutions (prepared in serum-free DMEM) to the appropriate column wells, bringing total well volumes to 100 μL. Columns 1 and 12 as well as the top and bottom rows of all 96-well plates were treated as mock wells and excluded from data analysis to control for plate edge effects (Burt et al. 1979). Immediately after adding 5X test solutions to plates, cells were returned to the incubator for the experiment duration (typically 3 days or approximately 72 hours) after which 20 μL of MTS reagent was added to each plate well. Plates were then incubated for another 4 hours. (For the experiment shown in Fig. 2d, only 10 μL of MTS reagent was added to plate wells because there was not enough reagent to add 20 μL to all 96 wells.) End-point absorbance measurements were taken with a plate reader at 490 nm (BioTek Instruments, Agilent Technologies).

### Cell Counting and Imaging

Cells were harvested, pelleted, and resuspended in serum-free DMEM. Two replicate 50 μL samples of cell suspensions were each diluted 1:1 in trypan blue dye, from which 10 μL was sampled and pipetted onto a hemocytometer (improved Neubauer ruling pattern). Live and dead/blue (non-intact) cell counts were performed in duplicate (i.e., one count per sample). Brightfield images were obtained using an inverted tissue culture microscope (Hund Wetzlar) fitted with a fixed microscope adapter connected to a 5.1-megapixel digital camera (AmScope). Original grayscale image tiles were merged into a single multi-panel image representing each experiment. Multi-panel images were processed with Fiji/ImageJ2 open-source software (NIH). Image processing involved pseudo flat field correction to balance uneven illumination across images, Fast Fourier Transform (FFT) bandpass filtering to reduce image scan lines, and brightness/contrast adjustments.

### Data Analyses and Statistics

LC-MS datasets, including both matrix LC and MS data from 3 replicate injections of the same sample type, were first averaged and normalized; mean data matrices were then visualized using R software (v4.1.0) with viridis and ggplot2 packages. Principal component analysis (PCA) on averaged MS data was performed in R using the prcomp function. Potential parent species were identified in a mass spectral database (MassBank.eu) by searching the *m*/*z* value and average relative intensity of peaks shared between samples. All other data were analyzed and/or visualized using Prism software (v9.2.0, GraphPad), in which two-sided *t* tests with Welch’s correction were used to determine statistical significance between compared groups.

## Results

### Molecular features of FBS and algae extracts show both distinct and common components

Given their potential growth effects, we sought to compare algae extracts (CVE and CGF, each of which was derived from a different *Chlorella* strain) to each other and FBS using LC-MS. Due to limited scope and the molecular complexity of these samples, our aim was not to look for specific compounds but rather to gain a qualitative profile for each sample type. Fig. 1a shows LC chromatograms of FBS compared to CVE and CGF samples. Large broad peaks around 260 nm for species that eluted off the column within the first few minutes were observed in both CVE and CGF but not FBS samples; these large peaks did not correspond to any appreciable signal(s) detected by the mass spectrometer at such early elution times (Fig. 1b). All samples showed both substantial absorbance (between 200-250 nm) and mass signals at retention times from approximately 18-21 minutes; the relative magnitude of this cluster of signals was comparable for FBS and CVE samples, whereas that of CGF samples was lower. Fig. 1c shows the results of a principal component analysis (PCA) comparing total ion chromatograms (TICs) of all FBS, CVE and CGF samples (i.e., replicates averaged in Fig. 1b). Replicate injections of the same sample type clustered together and separated in PC1 but not PC2 values; moreover, CVE and FBS samples were more similar in PC4 values than CGF samples. We also visualized how signals at each *m*/*z* value and retention time compared across samples (Fig. 1d). Each sample type showed a common pattern with a cluster of relatively high signal peaks corresponding to *m*/*z* values ranging from roughly 800-1200 and retention times from about 15-21 minutes. Both CVE and CGF samples also shared signal features at lower *m*/*z* values (mainly over the 400-800 range) and retention times from about 5-15 minutes; this broad cluster of signals was less apparent if not absent in FBS samples. We further examined the most striking MS feature found in all samples, i.e., the cluster of signals at *m*/*z* values from about 800-1200 and retention times from 15-21 minutes. Fig. 1e highlights five relatively abundant and shared peaks within this cluster, noting their *m*/*z* whole integer value, relative abundance, and corresponding potential parent ion identity(s). The top candidate identified by a mass spectral database search (MassBank.eu) for the most abundant shared peak (no. 3 in Fig. 1e) was nicotinamide adenine dinucleotide (NAD^+^) and/or NAD phosphate (NADP^+^).

**Fig. 1.**
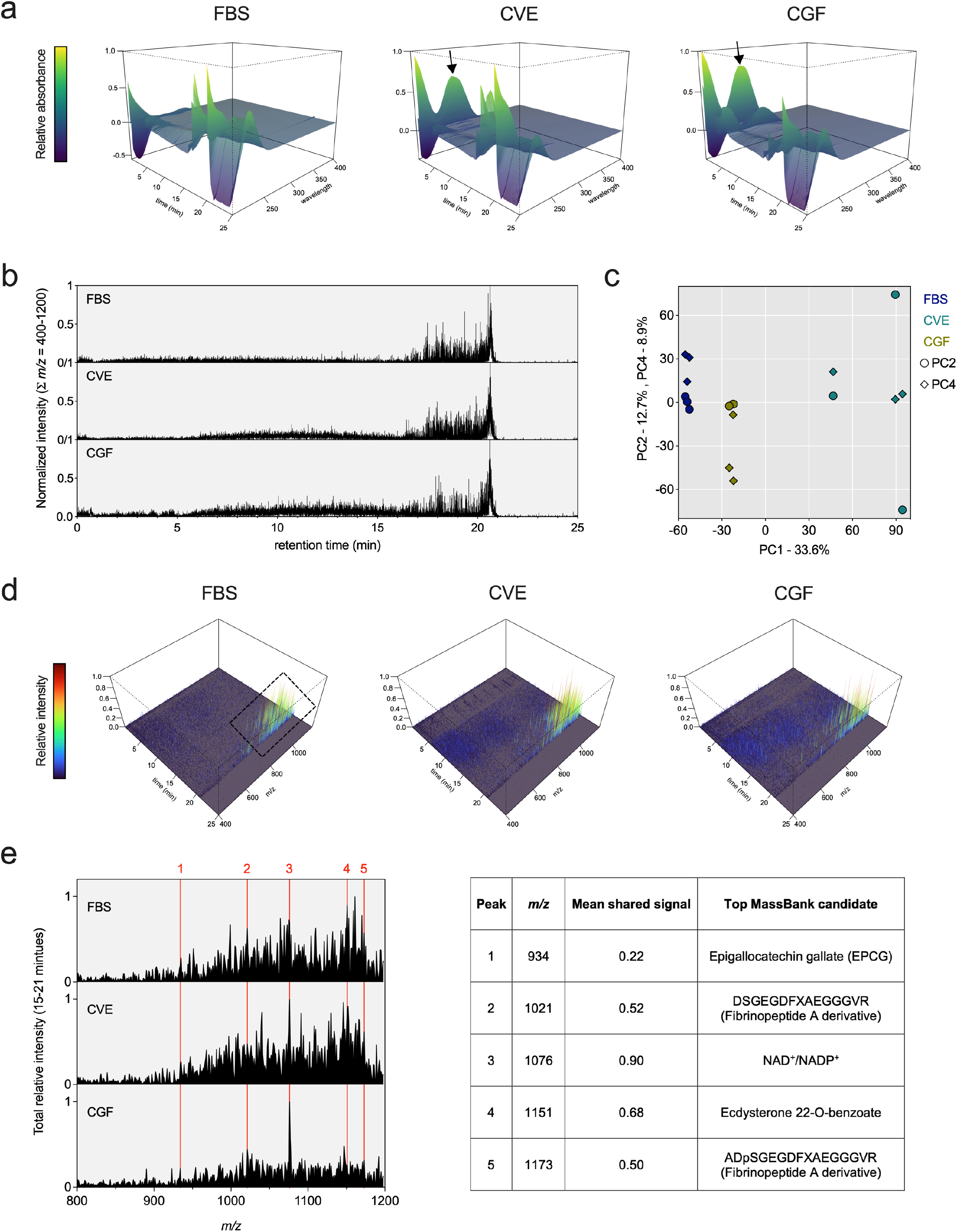
Comparative characterization of *Chlorella* extracts versus FBS by LC-MS. (**a**) Perspective 3D surface plots are shown comparing UV chromatograms of FBS, CVE and CGF samples (wavelengths 200-400 nm); each plot shows the mean normalized absorbance spectra of three replicate injections of the same sample type. Arrows indicate large, broad peaks around 260 nm present in CVE and CGF samples. (**b**) Normalized total ion chromatograms (TICs) are shown as the sum of signals of all *m*/*z* values from 400-1200 for FBS, CVE and CGF samples; representative data shown are averaged spectra from three injections of the same sample type. (**c**) Scatter plot shows results of principal component analysis (PCA) on panel b MS TIC data; sample types are indicated by different colors with PC2 (circles) and PC4 (diamonds) values plotted on the same y-axis scale. (**d**) Perspective 3D surface plots show normalized mass spectra of FBS, CVE and CGF samples; data shown are averages of three replicate injections. Dashed box indicates region of interest where signal location/intensity appears similar across all three sample types (i.e., retention times from 15-21 minutes and *m*/*z* values from 800-1200). (**e**) Aligned mass spectra show the sum of all signals for retention times from 15-21 minutes at each whole integer *m*/*z* value from 800-1200; data are shown as mean normalized spectra of three replicate injections of the same sample. Red vertical lines indicate signals of relatively high abundance shared across all three sample types; table lists potential parent species for corresponding signals based on top candidates identified by database (MassBank.eu) search

### Algae extracts alone in the absence of FBS slightly improve cell viability but not growth

We initially set out to test if algae extracts have growth effects on cells cultured in the absence of any FBS, using MTS assays as an initial readout and proxy for potential cell growth. Fig. 2 shows the relative viability of EBTr cells cultured over a 10,000-fold range of concentrations of CVE or CGF. Media supplemented with CVE and CGF alone significantly increased EBTr cell viability compared to serum-free controls; such effects were more pronounced for CVE than CGF and for relatively high concentrations of CVE. However, increases in relative cell viability due to CVE supplementation alone did not exceed beyond roughly 15% in any condition; moreover, and importantly, upon examining CVE-treated cells under the microscope immediately prior to performing MTS assays, we did not observe any convincing level of growth or cell doubling(s).

**Fig. 2.**
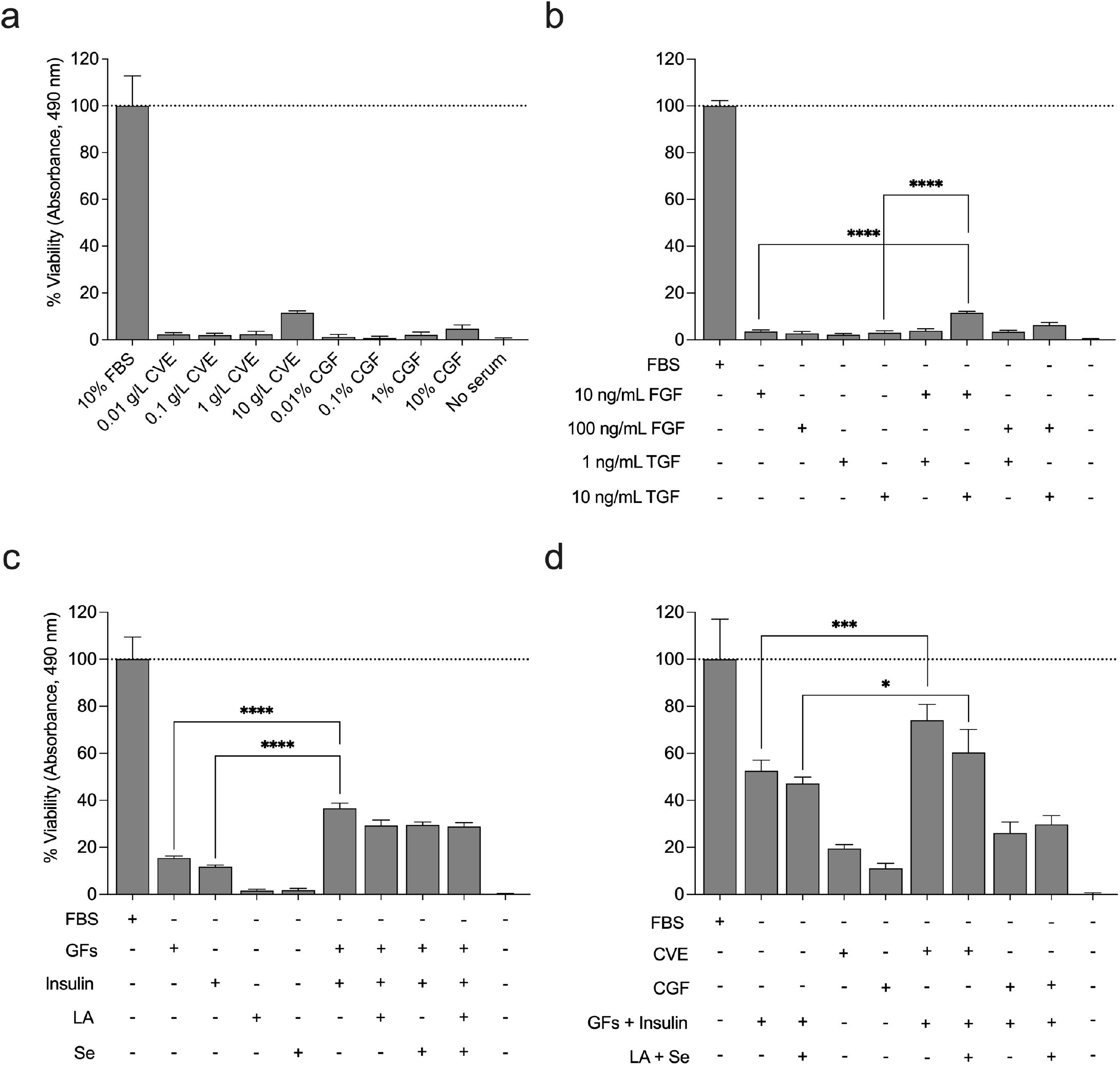
Select growth factor and insulin supplemented *C. vulgaris* extract improves cell viability in the absence of FBS. (**a**) Bar plots show viability of EBTr cells cultured with CVE and CGF relative to controls cultured in 10% FBS at day 3. Panel (**b**) shows relative viability of EBTr cells cultured in serum-free media supplemented with 10 or 100 ng/mL fibroblast growth factor 2 (FGF), or 1 or 10 ng/mL transforming growth factor 1 (TGF}, or combinations thereof, relative to controls cultured in 10% FBS after 3 days. Panel (**c**) shows relative viability of EBTr cells cultured in serum-free media supplemented with growth factors (GFs), or 3 μM bovine insulin, or 0.2 Mm L-ascorbate (LA), or 80 nM selenium dioxide (Se), or combinations thereof, relative to controls cultured in 10% FBS. Growth factors were used at 30 ng/mL (FGF) and 10 ng/mL (TGF). Panel (**d**) shows relative viability of EBTr cells cultured in serum-free media supplemented with 10 g/L CVE or 10% CGF alone, or GFs and insulin with and without LA and Se, or combinations thereof, compared to controls cultured in 10% FBS. Concentrations of GFs, insulin, LA and Se were the same as that for panel c. Statistical significance was determined by *t*-test with Welch’s correction; significance levels are indicated as \**p*<0.05, \*\*\**p*<0.001, \*\*\*\**p*<0.0001, ns=not significant, n=6

Given the partial but limited growth shown in Fig. 2a, we wondered if there may be components in the CVE which were acting to inhibit cell growth. We decided to separate the different components of CVE through filtration, centrifugation, and treatment with nuclease, using MTS assays to compare head-to-head the extract in its crude, filtered, supernatant, and pellet forms, each with and without nuclease treatment. Fig. S1a shows the relative viability of EBTr cells cultured with these different CVE fractions. No obvious differences were seen between crude, filtered and supernatant forms of extract, with viabilities of such fractions all falling near the range of 15-20%. The viability of cells treated with the resuspended pellet was notably lower than that of all other fractions, falling to nearly 0%, similar to that of no serum controls. Further, CVE fractions treated with nuclease showed mostly reduced viability compared to the corresponding fractions not treated with nuclease.

Fig. S1 (panels b and c) show the LC-MS characterization of the supernatant and pellet forms of CVE used in the experiment described in Fig. S1a. UV absorbance data shows sharp peaks around 20 minutes in the 200-250 nm range for both the pellet and the supernatant. The supernatant also shows a pair of broader peaks at around 200 and 260 nm that appear to have eluted within the first minute, neither of which was present in the pellet form. TICs of mass spectrometry data for *m*/*z* values between 400 and 1200 for these two forms of extract show similar footprints. Both show similarly the highest intensity spectra between 15 and 20 minutes, with relatively smaller spectra between 0 and 15 minutes. The spectra of the pellet form show slightly greater relative signals in the 0-6-minute range. Notably, the large UV peaks in both sample types correspond to increased signals in their respective mass spectra at around 20-21 minutes, however the large UV signal eluting early and only in the supernatant did not correspond to any appreciably large signal(s) in its mass spectra.

### Select serum-free media supplemented with *C. vulgaris* extract enhances cell viability

After seeing limited results from algae extract alone, we decided to move our research in a different direction, focusing on growth factors, which have previously been used to successfully culture stem cells in an “Essential 8” (E8™) serum-free medium (Chen et al. 2011). Fig. 2b shows percent viability of EBTr cells treated with different concentrations of basic fibroblast growth factor (FGF) and transforming growth factor β1 (TGF). Neither growth factor alone improved viability beyond 5-10% relative to serum-free controls, yet viability increased to around 15% for EBTr cells treated with a combination of 10 ng/mL FGF and 10 ng/mL TGF. Thus, we decided to use a combination of FGF and TGF for our next experiment in which we tested these growth factors together with bovine insulin, vitamin C (in the form of L-ascorbate), and selenium (in the form of selenium dioxide), all of which are listed as ingredients of the E8™ media. Fig. 2c shows insulin as a critical factor in real cell growth, in conjunction with growth factors; treatments with both insulin and growth factors improved EBTr cell viability to around 35% relative to controls cultured with FBS. Despite their inclusion in E8™ media, adding L-ascorbate, selenium, or both, slightly reduced EBTr cell viability compared to that of cells cultured with growth factors and insulin alone. We decided next to bring algae extracts back into our experiments, combining them with insulin and growth factors and/or L-ascorbate and selenium. Fig. 2d shows that EBTr cell viability reached approximately 75% (relative to FBS controls) when treated with a combination of CVE, growth factors and insulin. On the other hand, the viability of cells treated with CGF, growth factors and insulin, was actually lower than that of cells treated with growth factors and insulin alone. Once again, the addition of L-ascorbate and selenium had little if any beneficial effect on cell viability.

### Select serum-free media supplemented with *C. vulgaris* extract promotes cell growth and expansion

After seeing promising viability results (i.e., Fig. 2d), we decided the next step was to test if EBTr cells would expand when treated with insulin, growth factors and CVE in a cell count experiment. Fig. 3a shows cell counts of CVE and CGF with supplements (i.e., growth factors, insulin, L-ascorbate, and selenium) compared to controls with and without serum (10% FBS). The number of EBTr cells increased in cultures with CVE or CGF plus supplements by day 2, showing definitive growth, yet still not as much growth as control cells cultured with 10% FBS. By day 4, the number of cells cultured with CGF plus supplements fell to a level near that of no serum controls, but cells cultured with CVE plus supplements continued growth to around half that of controls cultured with 10% FBS. On day 8, the number of cells cultured with CVE plus supplements was comparable to that of controls cultured with 10% FBS, the latter of which had decreased since day 4. Fig. 3b shows the corresponding cell death percentages for the experiment described in Fig. 3a. Cell death percentages were largely in the 10-15% range, slightly increasing over time, with the most death occurring for cells cultured with CGF plus supplements on day 4 jumping to about 40%, corroborating the loss of growth shown for such cultures between day 2 and day 4 shown in Fig. 3a. Fig. 3c shows images captured of the experiment described in Fig. 3a and 3b. EBTr cells cultured with 10% FBS show sustained growth through day 4 until reaching confluency, at which point growth plateaued between days 4 and 8. Cells cultured with CVE plus supplements also appeared to grow through day 4, reaching a relatively high level of confluency; on day 8 the level of confluency appears about the same as day 4 (perhaps slightly more) with more dead cells in the image. Cells cultured with CGF plus supplements appear to show growth until day 2, though notably less than cells cultured with CVE plus supplements; however, by day 4 most if not all remaining cells appear dead, and by day 8, only dead cells remain in the image. (Note that EBTr cells in wells treated with algae extract appear larger than cells in wells where algae extract was not present.) The no serum control shows some cells with normal morphology on day 2, with many dead cells in the image as well; by days 4 and 8, hardly any cells remain besides a few dead cells.

**Fig. 3.**
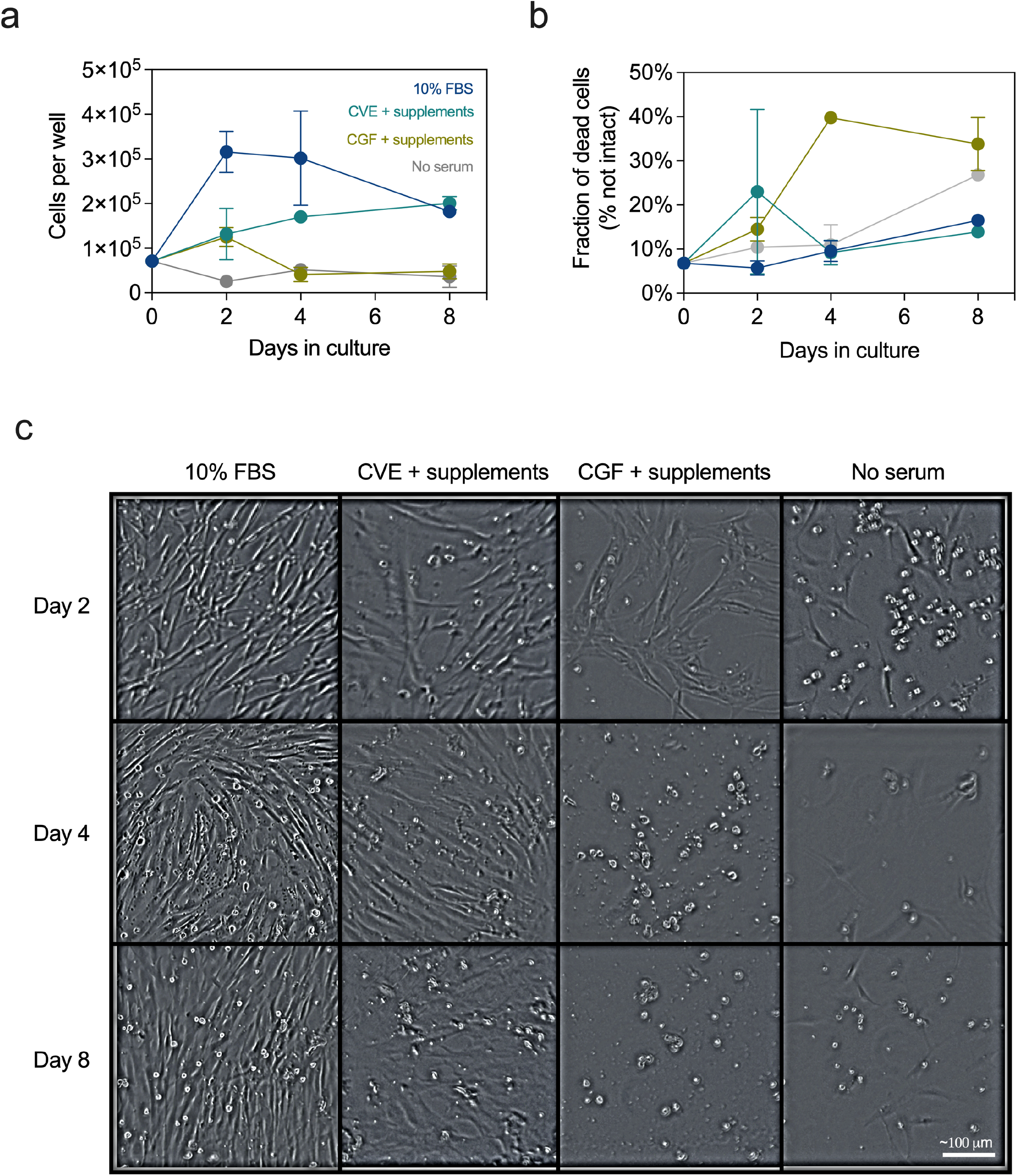
Growth factor and insulin supplemented *C. vulgaris* extract stimulates cell growth in the absence of FBS. Line plots show live (**a**) and percent dead/non-intact (**b**) EBTr cells after 0, 2, 4 and 8 days in culture. Different media conditions are indicated by different colors; all counts were performed in duplicate. (**c**) Brightfield images show representative EBTr cell density and morphology for each condition and time point; white scale bar indicates length of ∼100 μm. CVE was used at 10 g/L and CGF at 10% by volume. Supplements include 10 ng/mL TGF, 30 ng/mL FGF, 3 μM bovine insulin, 0.2 mM L-ascorbate, and 80 nM selenium

Given that EBTr cells showed growth and markedly improved viability when cultured with CVE and select supplements, we decided next to attempt to expand EBTr cells from a 1 mL culture (12-well plate) to a 2 mL culture (6-well plate) under these conditions (Fig. S2). On day 1, cells cultured with 10% FBS or with CVE plus supplements attached effectively, while the no serum control already shows many dead cells. By day 4, the no serum control well looks nearly identical to that on day 1, with some cells still attached but many dead with no visible growth. By day 4, cells cultured with 10% FBS or CVE plus supplements show obvious growth, nearing confluency. As observed in Fig. 3c, cells cultured with CVE plus supplements appear slightly bigger than cells not cultured with CVE, meaning that fewer cells were necessary to reach similar levels of confluency. After the day 4 pictures were taken, cells were detached and removed from the 1 mL cultures, then pelleted and resuspended into fresh 2 mL cultures. The following day, after cells attached to the plate (day 5), the relative amount of cells in the 10% FBS control well was notably greater than that in the well with CVE plus supplements. By day 8, cells cultured with 10% FBS reached near-confluency, while cells cultured with CVE plus supplements remained at nearly 50% confluency. On both days 5 and 8, the no serum control wells were essentially devoid of live cells, with only a few dead cells remaining.

## Discussion

Regarding our characterization of algae extracts compared to FBS, the most striking and obvious feature of the CVE and CGF samples that was not present in FBS samples were large absorbance peaks around 260 nm, which eluted off the column with the aqueous mobile phase within the first couple minutes. Such an absorbance profile is a characteristic property of nucleic acid (Setlow and Boyce 1960), and not surprising as a relatively abundant component present in crude cellular extracts. Moreover, since no appreciable mass signal(s) at this retention period was/were observed, it supports the notion that the species producing this absorbance spectra are likely nucleic acid species (e.g., DNA/RNA fragments containing at least 4 bases) with *m*/*z* values greater than that which could be detected (i.e., >1200). However, because the sensing of foreign nucleic acid often triggers cell death responses (Schlee and Hartmann 2016), we did not expect that foreign nucleic acid would be beneficial to cell growth/viability, rather on the contrary. Yet our results suggest that this putative nucleic acid at least did not harm cells and may have even been beneficial because nuclease-treated extracts, as well as the insoluble pellet with presumably very little nucleic acid (as evidenced by loss of the characteristic absorbance peak at 260 nm in Fig. S2a), both decreased cell viability as compared to cells given extract that was not pre-treated with nuclease or cells treated with soluble supernatant.

Despite an increase in EBTr cell viability observed for cells cultured with different concentrations of CVE and GCF alone, when such plates were observed under the microscope, no convincing amount of cell division was visible. When algae extract (CVE) was combined with growth factors, insulin and other additives, a similar marginal increase in cell viability of around 20% was observed in an environment where cells were already dividing. These observations of an increase in viability caused by CVE in both diving and non-dividing environments are consistent with other studies (Okamoto et al. 2022), suggesting that CVE increases cell viability independent of effects on proliferation. The possibility that algae extract enhances the cellular metabolic potential without promoting cell division requires further investigation. Cell viability assays using tetrazolium salts like MTS are often used as a proxy for cell growth, yet what such assays actually measure is the activity of mitochondrial NADH and NADPH-dependent dehydrogenase enzymes which reduce the reagent to a formazan product (Dunigan et al. 1995). In most cases, the more cells there are the more mitochondria are available to reduce the MTS reagent, but it is possible that the algae extracts have a positive effect on such metabolic activity without promoting cell growth. Given that a common and relatively abundant signal feature found in the mass spectra of algae extract samples mapped to NAD^+^/NADP^+^ as the parent species, it is possible that enriched levels of NAD(H)/NADP(H) present in the algae extracts may have caused enhanced activity of the dehydrogenase enzymes in the cells thus explaining the observed increase in viability. Further, it was previously shown that CVE also contains high levels of glutamate (Okamoto et al. 2020), as *Chlorella* algae use NADPH-dependent glutamate dehydrogenase enzymes to control their cell cycle (Talley et al. 1972). Glutamate has been shown to increase oxidation of NADH (Kannurpatti and Joshi 1999; Maalouf et al. 2007), and increase cancer cell viability and growth (Reiman et al. 1981; Takano et al. 2001; Gelb et al. 2015; Yamaguchi et al. 2020). Moreover, in fibroblasts, glutamate reduced toxicity induced by high levels of cysteine (Bannai and Ishii 1982), and glutamate dehydrogenase deficiency reduced cell viability (Tatsumi et al. 1989). Therefore, it is possible that enriched levels of glutamate in algae extracts may also be responsible for the increased viability observed.

Our experiments with components of the E8™ media clearly showed that insulin, basic-FGF and TGF-β are necessary for cell growth, and that these components are roughly additive in terms of their benefits to cell viability. Early studies examining serum-free culture conditions showed FGF and insulin as key determinants for growth (Mcfarland et al. 1991). Insulin causes cells to import glucose, providing energy and nucleotide building blocks needed for replication. In viability assays, addition of CVE also proved to be roughly additive, with CVE in addition to basic-FGF, TGF-β, insulin, and other additives combined yielding the highest viability reading of any serum-free condition tested. We demonstrated that EBTr cells can grow and expand into a new culture vessel when cultured in serum-free media that included at minimum basic-FGF, TGF-β, insulin and algae extract in the form of CVE, although such growth was not as robust as that of cells cultured with 10% FBS. Recent studies have shown that mitochondrial NADP(H) levels are essential for cell growth (Tran et al. 2021), without which cells cannot make proline required to support nucleotide and protein synthesis. Thus, it is plausible that enriched levels of NAD(H)/NADP(H) in CVE extract may provide a true benefit to cell growth; however, whether the additive increase in viability from CVE translates to any real benefit in growth rate beyond that from basic-FGF, TGF-β and insulin alone remains unclear. Regardless of whether algae extract is truly beneficial to cell growth, it is important to note that bovine fibroblasts could be cultured with only a subset of the E8™ components present. This may at least suggest some potential economic benefit due to lower cost serum-free formulations requiring fewer components. Since growth factors and insulin directly interact with their respective receptors through a relatively small range of residues across the entire protein, further experiments could test whether smaller peptide fragments mapping to such regions within these proteins remain actionable enough to sustain cell growth. Such substitutes may simplify and further lower the cost of serum-free formulations in which cells can be effectively cultured. Similarly, small molecules that act as agonists on basic-FGF and/or TGF-β receptors could also be tested for the same purpose.

Some limitations of our studies include aspects of the growth experiments in which we did not anticipate how differences in cell size could impact the results. In these experiments, it appears from the images as though cells cultured with algae extract were larger in size as compared to those not cultured with algae extract. Notably, increases in cell viability correlate with increased cell size (Dreesen and Fussenegger 2011), which could at least partially explain this observation since we also found that algae extract typically increased cell viability. However, differences in cell size observed with and without algae extract in the culture make the growth and expansion experiments difficult to interpret. For instance, during the expansion experiment, it appears that more cells were transferred from the 1 mL culture with 10% FBS compared to the culture with CVE plus supplements despite these two groups appearing to have a similar level of confluency on day 4. While both conditions promoted growth, the cells cultured in 10% FBS not only grow faster but require less space per cell. As a result, in the 2 mL cultures, the cells cultured in 10% FBS had a higher base count from which to start growing and were thus much more confluent by day 8. Despite the observed increase in viability, it is also a possibility that this increase in cell size corresponds to replicative senescence. Cells increase in size when they enter a permanent cell-cycle arrest and senescence (Biran et al. 2017), so further experiments examining the extent to which proliferation is affected by CVE are necessary. In a separate case, it should be noted that the day 8 FBS control result in our cell count experiment (Fig. 3c) shows a sharp decrease in cell number, which can be explained by over-confluence of the culture dish, resulting in cell death. As a result, the day 4 image and cell count more accurately display the efficacy of CVE plus supplements, since it would be expected that in a larger dish, the FBS control condition would continue to promote cell growth through day 8. Furthermore, we must acknowledge limitations in counting the fraction of dead cells recorded in our cell count experiments. As noted in the methods section, old media was first removed and the cells were briefly washed, which likely removed a large amount of previously dead and floating cells. Thus, the dead cell count more accurately represents the percentage of cells which were nearly dead, or dead, but not yet detached and floating. Finally, it is important to note that algae extracts may be subject to batch-to-batch variation depending on the growth and extraction conditions used to generate them, thus further study of such variation in *C. vulgaris* extract is important to confirm the reproducibility of results.

Taken together, our studies show that algae extract derived from *Chlorella vulgaris* enhances viability of bovine fibroblasts, possibly due at least in part to its enriched levels of NAD(H)/NADP(H) and/or glutamate, and that CVE can be used together with other supplements (basic-FGF, TGF-β, and insulin) to grow cells in the absence of serum. Further studies are needed to determine the extent to which CVE promotes cell growth and whether the supplementary components can be further modified to reduce their cost.

## Acknowledgements

This work was supported by the Figge-Bourquin Award and by Student-Faculty Collaborative Grants via the Creativity and Innovation and Dean’s Offices at Colorado College. The authors thank Rachel Wonciar and Tanya Cervantes in the Department of Chemistry and Biochemistry, and Carrie Moon and Darrell Killian in the Department of Molecular Biology at Colorado College, for their technical assistance.

## Author Contributions

G.D.C. and T.M.G. designed the study, performed experiments, analyzed data, and wrote the paper, with both authors contributing equally.

## Statements and Declarations

There are no competing interests.

