## Supplementary Material for "In vitro culture of bovine fibroblasts using select serum-free media supplemented with *Chlorella vulgaris* extract"

**Fig. S1**

**Nuclease-treated *C. vulgaris* extract and its insoluble fraction decrease cell viability. (a)** Bar plots shows relative viability of EBTr cells cultured with different fractions of 10 g/L CVE with and without nuclease treatment relative to control cells cultured in 10% FBS after 3 days. Statistical significance was calculated by *t*-tests with Welch's correction in which experimental CVE fractionation groups were compared with versus without nuclease treatment; significance levels are indicated as *$p<0.05$, ***$p<0.001$, ns=not significant, n=6. **(b)** Perspective 3D surface plots show UV chromatograms of CVE supernatant and pellet fractions (wavelengths 200-400 nm); data shown are mean normalized absorbance spectra from three replicate injections of each sample type. Arrow indicates large, broad peak around 260 nm present in CVE supernatant. Panel **(c)** shows normalized total ion chromatograms (TICs) for the sum of signals of all $m/z$ values from 400-1200 for CVE pellet and supernatant; representative data shown are averaged spectra from three injections of the same sample type.

**Fig. S2**

**Bovine fibroblasts cultured with select serum-free media and *C. vulgaris* extract show growth and expansion.** Brightfield images show representative EBTr cell density and morphology for each condition and time point. CVE was used at 10 g/L; supplements include 10 ng/mL TGF, 30 ng/mL FGF, and 3 μM bovine insulin. Cells were passaged and expanded from 1 to 2 mL cultures (and from 12-well to 6-well plates) immediately after day 4 images were taken.

Figure S1

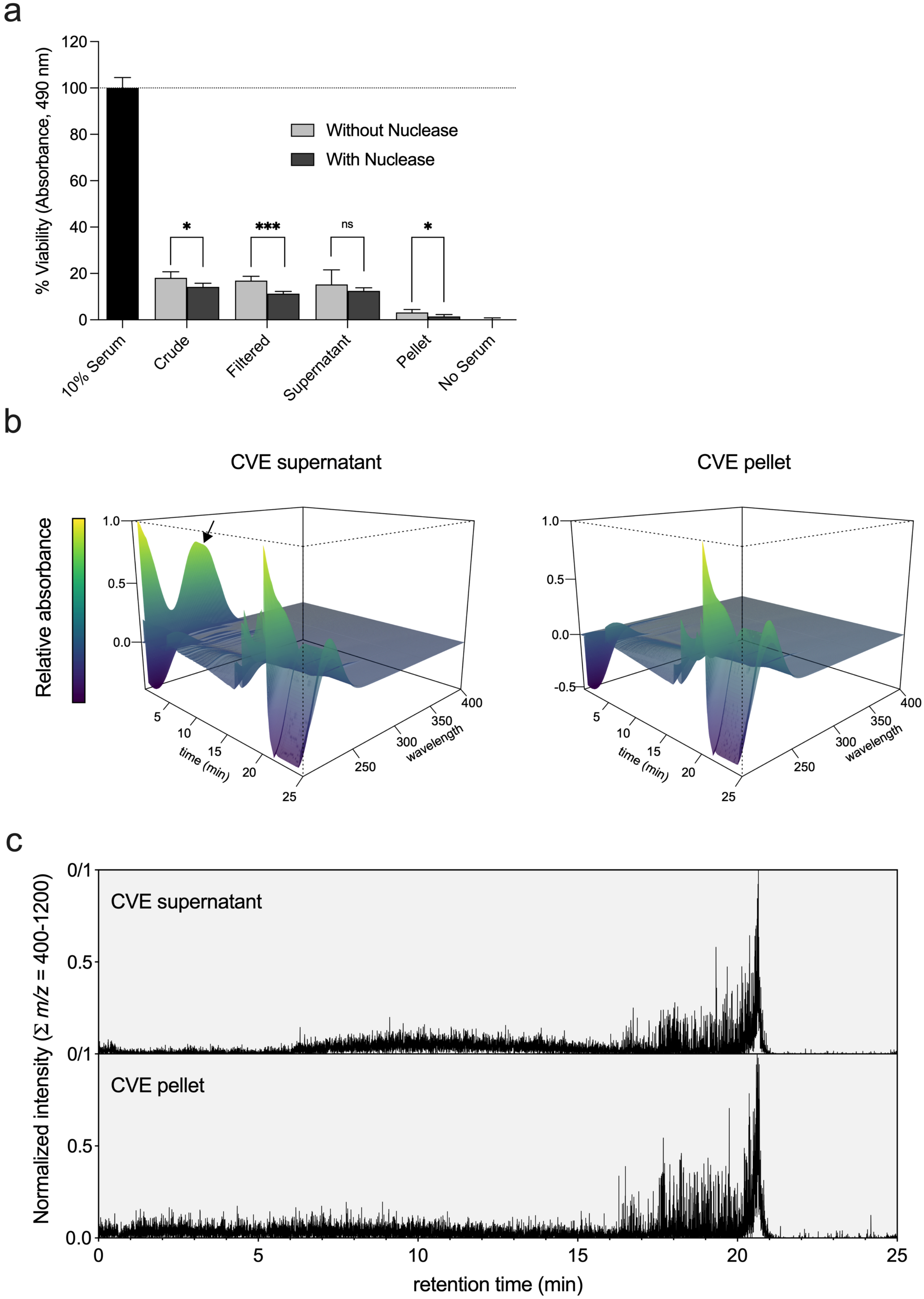

a
% Viability (Absorbance, 490 nm)
Without Nuclease
With Nuclease
*
***
ns
*
10% Serum
Crude
Filtered
Supernatant
Pellet
No Serum
b
CVE supernatant
CVE pellet
Relative absorbance
time (min)
wavelength
c
CVE supernatant
CVE pellet
Normalized intensity (Σ m/z = 400-1200)
retention time (min)

Figure S2

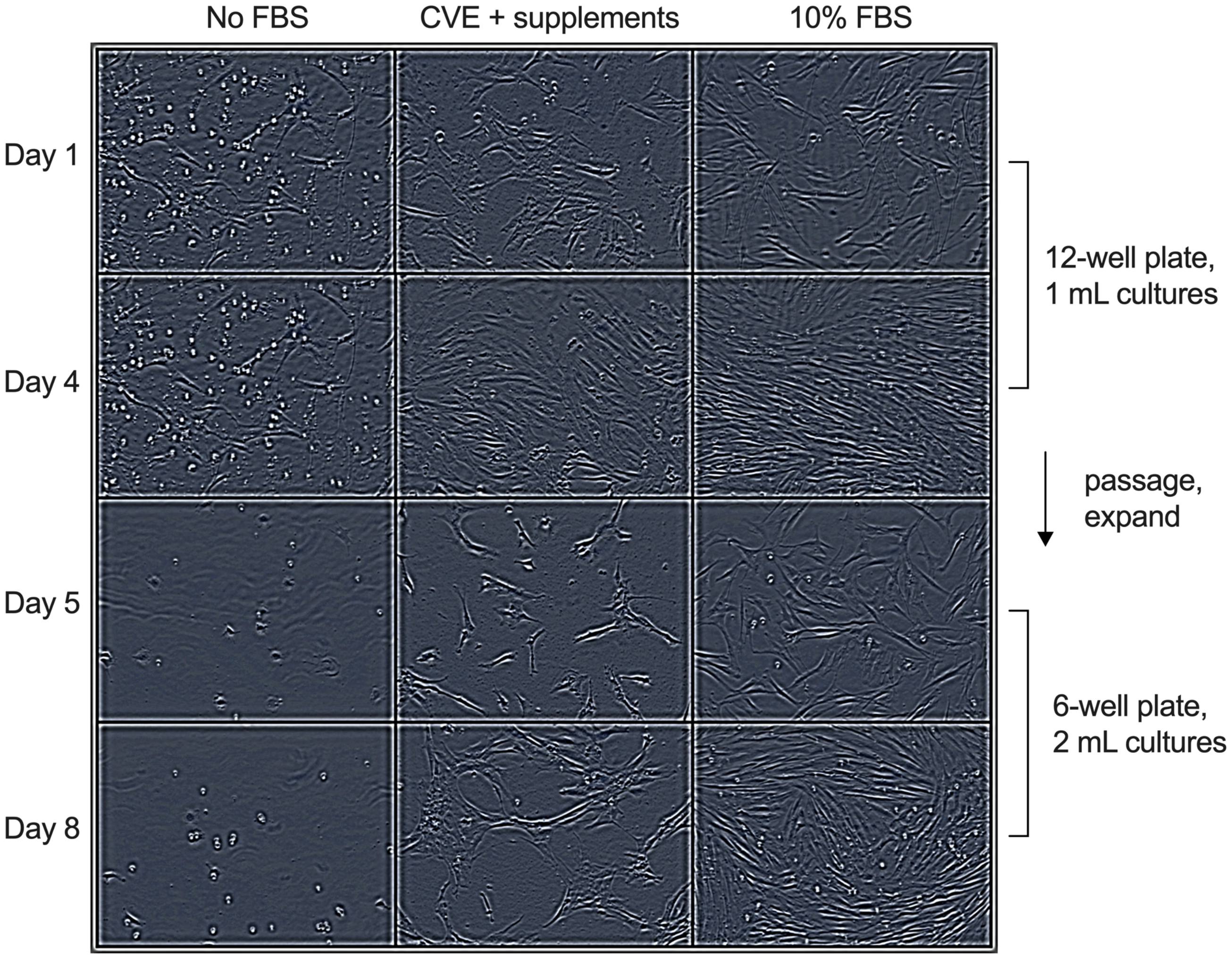

No FBS
CVE + supplements
10% FBS
Day 1
Day 4
Day 5
Day 8
12-well plate,
1 mL cultures
passage,
expand
6-well plate,
2 mL cultures
